# A chromosome-scale genome assembly of Timorese crabgrass (*Digitaria radicosa*): a useful genomic resource for the Poaceae

**DOI:** 10.1101/2024.05.14.594087

**Authors:** Koki Minoji, Toshiyuki Sakai

**Author notes:** Corresponding author: Toshiyuki Sakai.

## Abstract

Timorese crabgrass (*Digitaria radicosa*) is a grass species commonly found in Southeast Asia and Oceania. *Digitaria* species have high intraspecific and interspecific genetic and phenotypic diversity, suggesting their potential usefulness as a genetic resource. However, as the only high-quality reference genome available is for a tetraploid *Digitaria* species, a reference genome of the diploid species *D. radicosa* would be a useful resource for genomic studies of *Digitaria* and Poaceae plants. Here, we present a chromosome-level genome assembly of *D. radicosa* and describe its genetic characteristics; we also illustrate its usefulness as a genomic resource for Poaceae. We constructed a 441.6 Mb draft assembly consisting of 61 scaffolds with an N50 scaffold length of 41.5 Mb, using PacBio HiFi long reads. Thirty scaffolds, encompassing 440.8 Mb, were anchored to nine pseudochromosomes. We predicted 26,577 protein-coding genes, reaching a BUSCO score of 96.5%. To demonstrate the usefulness of the *D. radicosa* reference genome, we investigated the evolution of *Digitaria* species and the genetic diversity of Japanese *Digitaria* plants based on our new reference genome. We also defined the syntenic blocks between *D. radicosa* and Poaceae crops and the diverse distribution of representative resistance genes in *D. radicosa*. The *D. radicosa* reference genome presented here should help elucidate the genetic relatedness of *Digitaria* species and the genetic diversity of *Digitaria* plants. In addition, the *D. radicosa* genome will be an important genomic resource for Poaceae genomics and crop breeding.

## Introduction

Timorese crabgrass (*Digitaria radicosa*) is a diploid (2n = 18) plant commonly found in Southeast Asia and Oceania. The genus *Digitaria* comprises over 250 species in the family Poaceae, distributed mostly in tropical and warm temperate regions around the world (POWO, 2024). *Digitaria* species are mainly considered to be agricultural weeds, with 17 species registered as invasive plant species in CABI 2024. Nonetheless, several species are useful resources for agriculture. For example, fonio (*Digitaria exilis*) is an African indigenous cereal crop with highly nutritious grains, drought tolerance, and adaptation to nutrient-poor soils (Ayenan *et al*. 2018). Iburu (*Digitaria iburua*) and raishan (*Digitaria compacta*) are two other indigenous grain crops (Jideani 1999; Prance and Nesbitt 2004). Pangola grass (*Digitaria eriantha*) and hairy crabgrass (*Digitaria sanguinalis*) are utilized for livestock pastures and to control soil erosion (Pitman *et al*. 2016; Weinert-Nelson *et al*. 2022). Thus, *Digitaria* species can be weeds or crops, underscoring the interspecific phenotypic diversity of this genus. Therefore, understanding the genetic relatedness of *Digitaria* species may help to elucidate the weediness of Poaceae species and improve the utilization of *Digitaria* species in agriculture.

*Digitaria* plants also show high intraspecific genotypic and phenotypic diversity (Tsuyuzaki 2005; Scarcelli *et al*. 2011; Abrouk *et al*. 2020; Fukano *et al*. 2020; Wang *et al*. 2021). An exploration of the relationship between phenotypes and genotypes of diversified *Digitaria* populations will help determine the genetic architecture of various Poaceae traits. However, only a single chromosome-scale reference genome for a *Digitaria* species is currently available, for the cultivated tetraploid *Digitaria* species *D. exilis* (Abrouk *et al*. 2020; Wang *et al*. 2021). The polyploidy of this reference genome and the limited number of reference genomes hamper efforts to perform genomic analyses of *Digitaria* species (Ravet *et al*. 2018; Gonçalves Netto *et al*. 2021). In addition, *Digitaria* species have previously been classified using morphology-based approaches or from few DNA markers (Vega *et al*. 2009; Soreng *et al*. 2022; Touafchia *et al*. 2023). Notably, these two classification systems do not agree; a consensus classification remains unclear. Therefore, a reference genome for a diploid *Digitaria* species will facilitate genomic analysis, including population genomics and evolutionary analysis, to understand the genetic relatedness of *Digitaria* species and the genetic contribution to phenotypic diversity (Ravet *et al*. 2018).

Blast disease caused by the fungus *Magnaporthe* spp. is devastating for Poaceae species including rice (*Oryza sativa*) and *Digitaria* (Ou 1980). *Digitaria* plants commonly grow alongside rice at the edge of paddy fields in Japan (**Figure S1**). Notably, there is host plant specificity, with blast fungi isolated from *Digitaria* being *Magnaporthe grisea*, while isolates from rice are *Magnaporthe oryzae* (Tosa and Chuma 2014). Host specificity is determined by the interaction between effector proteins secreted by the fungus and resistance genes from the host plant (Tosa and Chuma 2014; Yoshida *et al*. 2016). Effectors secreted by *M. oryzae* are recognized by nucleotide-binding domain and leucine-rich repeat (NLR) proteins, which activates the resistance response (Cesari *et al*. 2013; Maqbool *et al*. 2015). *Digitaria* species inhabiting the same niche as rice plants may therefore have NLRs that recognize effectors secreted by *M. oryzae* isolates. Genomic resources for Japanese *Digitaria* species may thus provide resistance genes that can be deployed in rice against *M. oryzae*.

In this study, we constructed a highly contiguous and high-quality reference genome for the diploid species *D. radicosa*. We performed an evolutionary analysis using sequencing data from multiple *Digitaria* species and a population genetic analysis using *D. radicosa* and *Digitaria ciliaris* collected in Japan, based on our new reference genome. We also identified syntenic blocks between *D. radicosa* and Poaceae crops and the distribution of NLR genes in *D. radicosa*. These results highlight the utility of this new chromosome-scale diploid *Digitaria* reference genome for Poaceae genomics.

## Materials and Methods

### Sample collection

To generate the reference genome, one *Digitaria radicosa* plant was collected from Mozume, Kyoto Prefecture, Japan, in August 2022 and kept in a growth chamber at 32°C with a 12/12-hr light/dark cycle. Ten *D. radicosa* plants were sampled for population genomics analysis: four from Kyoto Prefecture, two from Kobe Prefecture, and four from Wakayama Prefecture, Japan (**Table S1**), in September 2023. Eleven *Digitaria ciliaris* plants were also sampled for population genomics analysis from paddy fields, coastal areas, and park areas in September 2023: five from Kyoto Prefecture and six from Wakayama Prefecture, Japan (**Figure S1, Table S1**).

### DNA isolation, library construction, and genome sequencing

For sequencing by HiFi reads, genomic DNA was extracted from one *D. radicosa* plant using 1 g of lyophilized leaf tissue and Qiagen Genomic-tips 20/G. The DNA was then size-selected using a Short Read Eliminator XS kit (PacBio) and g-TUBEs (Covaris). HiFi libraries were constructed using a SMRTbell Express Template Prep Kit 2.0 (PacBio); the resulting HiFi libraries were sequenced on a PacBio Revio instrument. For short-read sequencing, genomic DNA was extracted from the same *D. radicosa* plant using 100 mg of lyophilized leaf tissue and a Maxwell RSC Genomic DNA Kit (Promega). Sequencing libraries were constructed using a MGIEasy PCR-Free DNA Library Prep Set (MGI); the resulting libraries were sequenced on a DNBSEQ-G400 instrument as paired-end 150-bp reads. For population genomic analysis, a MIG-seq library was constructed. Genomic DNA was extracted from *D. radicosa* and *D. ciliaris* plants using 20 mg of young leaf tissue with a DNeasy Plant Mini Kit (Qiagen). After the DNA concentrations were adjusted to 10 ng/μL, the first PCR was performed using a Multiplex PCR Assay Kit Ver.2 (Takara) with MIG-seq primer set-1 developed by Suyama and Matsuki (2015). The PCR program followed Nishimura *et al*. (2022) with the exception of the initial denaturation, performed at 94°C for 1 min. The first PCR products were diluted 50-fold in ultrapure water. Indexing primers were added to each sample during the second PCR, which was performed with PrimeSTAR GXL DNA Polymerase (Takara) following the PCR program of Nishimura *et al*. (2022) but with a final extension of 7 min at 68°C. The concentration of each amplicon was measured and adjusted before pooling equal amounts of DNA from each sample. The pooled library was purified using AMPure XP Beads (Beckman Coulter). The library was then size-selected using SPRIselect (Beckman Coulter). The bead ratios for the left size selection and right size selection were set to 0.75× and 0.56×, respectively. The library was sequenced on an Illumina NovaSeq platform as paired-end 150-bp reads by Nippon Genetics Co., Ltd.

### Genome assembly and evaluation

Genome size was estimated within the MaSuRCA pipeline based on the k-mer abundance distribution using the short reads generated on the DNBSEQ instrument (Zimin *et al*. 2013). Primary contigs were assembled using PacBio HiFi reads by the hifiasm v0.19.8 assembler (Cheng *et al*. 2021). The primary genome assembly was polished using HiFi reads and short reads by NextPolish2 v0.2.0 (Hu *et al*. 2024). To remove redundant sequences, purge_dups v1.2.5 was used (Guan *et al*. 2020). The assembled contigs were then scaffolded into pseudochromosomes by mapping against the chromosome-scale *D. exilis* genome, using Ragtag v2.1.0 (Abrouk *et al*. 2020; Alonge *et al*. 2022). To assess completeness of the assembled genome, a Benchmarking Universal Single-Copy Orthologs (BUSCO) v5.6.1 analysis was performed with the poales_odb10 dataset (Manni *et al*. 2021). All scripts used to perform genome assembly and evaluation were deposited in a GitHub repository (https://github.com/slt666666/D_radicosa_genome).

### Genome annotation

The *D. radicosa* genome was annotated using the RepeatModeler2 pipeline, which identifies repeats in the genome, and the BRAKER3 pipeline for *de novo* gene prediction, homology-based predictions, and transcriptome-based predictions to predict protein-coding genes (Flynn *et al*. 2020; Gabriel *et al*. 2023). For the annotation of repeat elements, RepeatModeler2 v2.0.5 was used to perform *ab initio* prediction with default parameters (Flynn *et al*. 2020). The annotated repeats were then filtered to only keep complex repeats for soft-masking. Repeat elements across the genome were masked before gene annotation by RepeatMasker v4.1.5 (Tarailo-Graovac and Chen 2009). Next, BRAKER3 was used to train GeneMark-ETP and AUGUSTUS on the soft-masked genome. For transcriptome-based gene annotation, 12 public transcriptome deep sequencing (RNA-seq) libraries generated for *Digitaria* species were downloaded from NCBI and mapped to the *D. radicosa* genome using STAR v2.7.10a to generate mapped BAM files (**Table S2**) (Dobin *et al*. 2013). For homology-based gene annotation, protein sequences from *Arabidopsis thaliana* (GCA_000001735.2), *Setaria italica* (GCA_000263155.2), *Digitaria exilis* (GCA_015342445.1), *Sorghum bicolor* (GCA_000003195.3), *Oryza sativa* (GCA_001433935.1), and *Zea mays* (GCF_902167145.1) were downloaded from NCBI. To validate the quality of predicted gene sets, completeness was assessed using BUSCO v5.6.1 with the poales_odb10 dataset (Manni *et al*. 2021). Completeness and lineage consistency of the proteome (the proportion of protein sequences placed into known gene families from the same lineage) were also assessed using OMArk (Nevers *et al*. 2024).

### Phylogenetic analysis

Publicly available sequencing reads for the genomes and transcriptomes of *Digitaria* species (*Digitaria abyssinica, Digitaria bicornis, Digitaria biformis, Digitaria brownii, Digitaria californica, Digitaria ciliaris, Digitaria ctenantha, Digitaria cuyabensis, Digitaria didactyla, Digitaria eriantha, Digitaria exilis, Digitaria fuscescens, Digitaria longiflora, Digitaria milanjiana, Digitaria monodactyla, Digitaria pseudodiagonalis, Digitaria sanguinalis*, and *Digitaria ternata*) were used for phylogenetic analysis. The accession numbers for all raw reads are listed in **Table S2**. The obtained DNA-seq and RNA-seq reads were filtered and trimmed using fastp (Chen *et al*. 2018). The quality-trimmed DNA-seq reads were mapped to the new *D. radicosa* reference genome assembled in this study using bwa-mem2 (Vasimuddin *et al*. 2019). RNA-seq reads were mapped using STAR v2.7.10a (Dobin *et al*. 2013). Variant calling was performed using GATK according to GATK Best Practices recommendations (McKenna *et al*. 2010). Single-nucleotide polymorphisms (SNPs) at biallelic sites with read depth >1 in all samples were used for phylogenetic analysis. A maximum-likelihood phylogenetic tree was estimated using RAxML (Stamatakis 2006). The GTRCAT model was selected as the best substitution model based on the Akaike information criterion calculated by ModelTest-NG (Darriba *et al*. 2020). All scripts and criteria of quality-filter used to perform the phylogenetic analysis were deposited in a GitHub repository (https://github.com/slt666666/D_radicosa_genome).

### Population genetics

The reads obtained from the MIG-seq library were used for population genetics analysis. The raw reads were trimmed and quality-filtered using fastp v0.23.2 (Chen *et al*. 2018). The trimmed reads were mapped to the new *D. radicosa* genome assembled in this study using BWA v0.7.17 (Li 2013). Alignments with mapping quality below 40 and reads with multiple alignments were removed using SAMtools v1.19.2 (Li *et al*. 2009). Each bam file containing reads from a single sample were combined to a single bam file, and variant calling was performed using bcftools v1.16 (Li 2011). Three Variant Call Format (VCF) files were created, containing SNPs from *D. radicosa, D. ciliaris*, or both species. Filtering of SNPs was performed on the number of alleles (only two-allele loci were retained), minor allele frequency (MAF ≥ 0.05), genotyping quality (GQ ≥ 20), and read depth (within a range of 5 to 50) using bcftools and VCFtools v0.1.17 (Danecek *et al*. 2011). SNPs within possible linkage disequilibrium regions were then pruned based on (1) physical distance between SNPs using the bcftools ‘+prune’ plugin with options ‘-n 1 -w 1kb’ and (2) genotypic correlation between SNPs using the PLINK version 1.90 ‘—indep-pairwise’ option with settings ‘5000kb 1 0.1’ (Chang *et al*. 2015). Using these filtered SNP datasets, a principal component analysis (PCA) was performed using the PLINK ‘—pca’ option. The result of PCA with all samples suggested misidentification of two samples as *D. ciliaris*, so these two samples were removed before repeating the analysis (**Figure S2**). All scripts, criteria of quality filters, and files necessary to reproduce the analysis were deposited in a GitHub repository (https://github.com/slt666666/D_radicosa_genome).

### Synteny analysis

MCScanX was used to define syntenic blocks between the *D. exilis* and *D. radicosa* genomes and between the *O. sativa* and *D. radicosa* genomes (Wang *et al*. 2012; Sakai *et al*. 2013; Abrouk *et al*. 2020). We used a rice reference genome of Nipponbare cultivar (IRGSP-1.0) (Sakai *et al*. 2013). The ‘-b 2’ option was used to identify patterns of syntenic blocks between two species. The orthologous gene set used for synteny analysis with MCScanX was identified by BLASTP search with an e-value ≤ 1e−10. At least 10 contiguous genes were required for synteny calling in each block. The results of MCScanX were visualized using SynVisio (Bandi *et al*. 2022).

### NLR annotation and phylogenetic analysis

NLRtracker was used to identify NLR genes from the new *D. radicosa* genome (Kourelis *et al*. 2021). These NLRs from *D. radicosa* and functionally validated NLRs from Poaceae species in the RefPlantNLR database were used for phylogenetic analysis (Kourelis *et al*. 2021). NLR protein sequences were aligned using MAFFT v.7, deleting the gaps in the alignments (Katoh and Standley 2013). Sequences of NLRs with truncated NB-ARC domains were manually removed. The sequences of the NB-ARC domains from the aligned NLR proteins were used to reconstruct phylogenetic trees. A maximum-likelihood phylogenetic tree was generated in RAxML version 8.2.12 using the JTT model. The bootstrap values in the resulting tree were based on 100 iterations (Stamatakis 2006).

## Results

### Genome assembly and genome annotation

We assembled a reference genome for Timorese crabgrass (*D. radicosa*, 2n = 18) using PacBio HiFi long reads and DNBSEQ short reads. To estimate the size of the *D. radicosa* genome, we analyzed 88.3 Gb of short reads based on the k-mer abundance distribution, revealing an estimated genome size of 420 Mb. To obtain the reference genome, we assembled 21.7 Gb of HiFi long reads and polished the resulting contigs with 88.3 Gb of DNBSEQ short reads. We obtained 61 contigs covering 441.6 Mb in total length with a contig N50 value of 41.5 Mb. The length of the longest contig was 66.2 Mb, indicating that some contigs were assembled at the scale of a chromosome. These assembly statistics reflect the good contiguity and integrity of our genome assembly (**Table 1**). To generate a chromosome-scale genome assembly, we performed reference-guided assembly using the *D. exilis* genome, after which the 30 longest contigs encompassing 440.8 Mb of sequence were anchored to nine pseudochromosomes (**Table 1**). The GC content of the genome was 45.4% (**Table 1**). To evaluate the quality of the assembled genome, we performed a Benchmarking Universal Single-Copy Orthologs (BUSCO) analysis using the Poales lineage gene dataset. We calculated a BUSCO score of 96.3% (4,714 out of 4,896) for the newly assembled genome, with 93.4% single-copy and 2.9% duplicated genes (**Table 1**). These results indicate that the chromosome-scale assembled *D. radicosa* genome in this study is highly contiguous and of high quality.

**Table 1.** Summary statistics for genome assembly and genome annotation.

| Item | Category | Number |
| --- | --- | --- |
| Sequencing | PacBio HiFi reads | 21.7 Gb |
|  | Illumina short reads | 88.3 Gb |
| Assembly | Estimated genome size | 420.3 Mb |
|  | Assembled genome size | 441.6 Mb |
|  | Contig number | 61 |
|  | Contig N50 | 41.5 Mb |
|  | Longest contig | 66.2 Mb |
|  | Scaffold number | 40 |
|  | Scaffold N50 | 51.8 Mb |
|  | Longest scaffold | 66.2 Mb |
|  | Genome anchored to chromosome | 99.8% |
|  | Complete BUSCOs (poales_odb10) | 96.3% (4,714/4,896) |
|  |  | S:93.4%, D:2.9%, F:0.3%, M:3.4%* |
| Annotation | GC content | 45.4% |
|  | Repeat sequences | 50.1% |
|  | Number of protein-coding genes | 26,577 |
|  | BUSCO score (poales_odb10) | 95.3% (4,665/4,896) |
|  |  | S:92.6%, D:2.7%, F:0.2%, M:4.5%* |

\*S, single; D, duplicated; F, fragmented; M, missing.

To predict protein-coding genes in the genome, we performed an *ab initio* prediction of repeat elements to mask them in the genome, after which we subjected the masked genome to our gene prediction pipeline. We identified 223.6 Mb of repetitive elements in the *D. radicosa* genome, representing 50.1% of the entire genome (0.1% of short interspersed nuclear elements [SINEs], 1.6% of long interspersed nuclear elements [LINEs], 16.5% of long terminal repeat elements, 3.7% of DNA transposons, and 28.3% of unclassified repeats) (**Table S3**). The gene prediction pipeline identified 26,577 protein-coding genes (Table 1). We assessed the completeness of the gene prediction by repeating the BUSCO analysis with the same Poales lineage gene dataset. We identified 95.3% (4,665 out of 4,896) complete BUSCO genes among the predicted protein-coding genes, with 92.6% as single copies and 2.7% as duplicated genes (**Table 1**). We also performed an analysis with the OMArk software, which returned a level of completeness of 90.6% (18,575 out of 20,501) and a taxonomic consistency of 93.2% (24,775 out of 26,577) (**Table S4**). These results indicate the high completeness and accuracy of our predicted gene set.

### Evolutionary analysis of *Digitaria* species

The current classification of *Digitaria* species is not consistent between studies due to different methodologies (Clayton *et al*. 2021; WCSP 2022). However, the issues of relatedness among *Digitaria* species should be solved by a genomic analysis based on high-quality genome data. To investigate the genetic relatedness of *Digitaria* species, we thus performed a phylogenetic analysis based on our new *D. radicosa* reference genome using publicly available DNA-seq and RNA-seq data for 22 samples from 19 *Digitaria* species (**Table S2**). We used *Chlorocalymma cryptacanthum* as an outgroup. We aligned the publicly available DNA-seq and RNA-seq reads to our reference genome and identified 2,052 SNPs with a read depth over 2 for all 23 samples. The resulting phylogenetic tree reconstructed from the obtained SNPs showed that *Digitaria* species can be classified into two well-supported clades (**Figure 1**). The first clade contains a cultivated *Digitaria* species, *D. exilis*, and its putative wild progenitor, Indian crabgrass (*D. longiflora*). The second clade contains Asian weedy *Digitaria* species including *D. ciliaris* and *D. radicosa*. While the close relationship of *D. exilis* and *D. longiflora* was known (Abrouk *et al*. 2020), we noticed that *D. fuscescens* is also phylogenetically closely related to *D. exilis* in our phylogenetic analysis (**Figure 1**).

**Figure 1.**
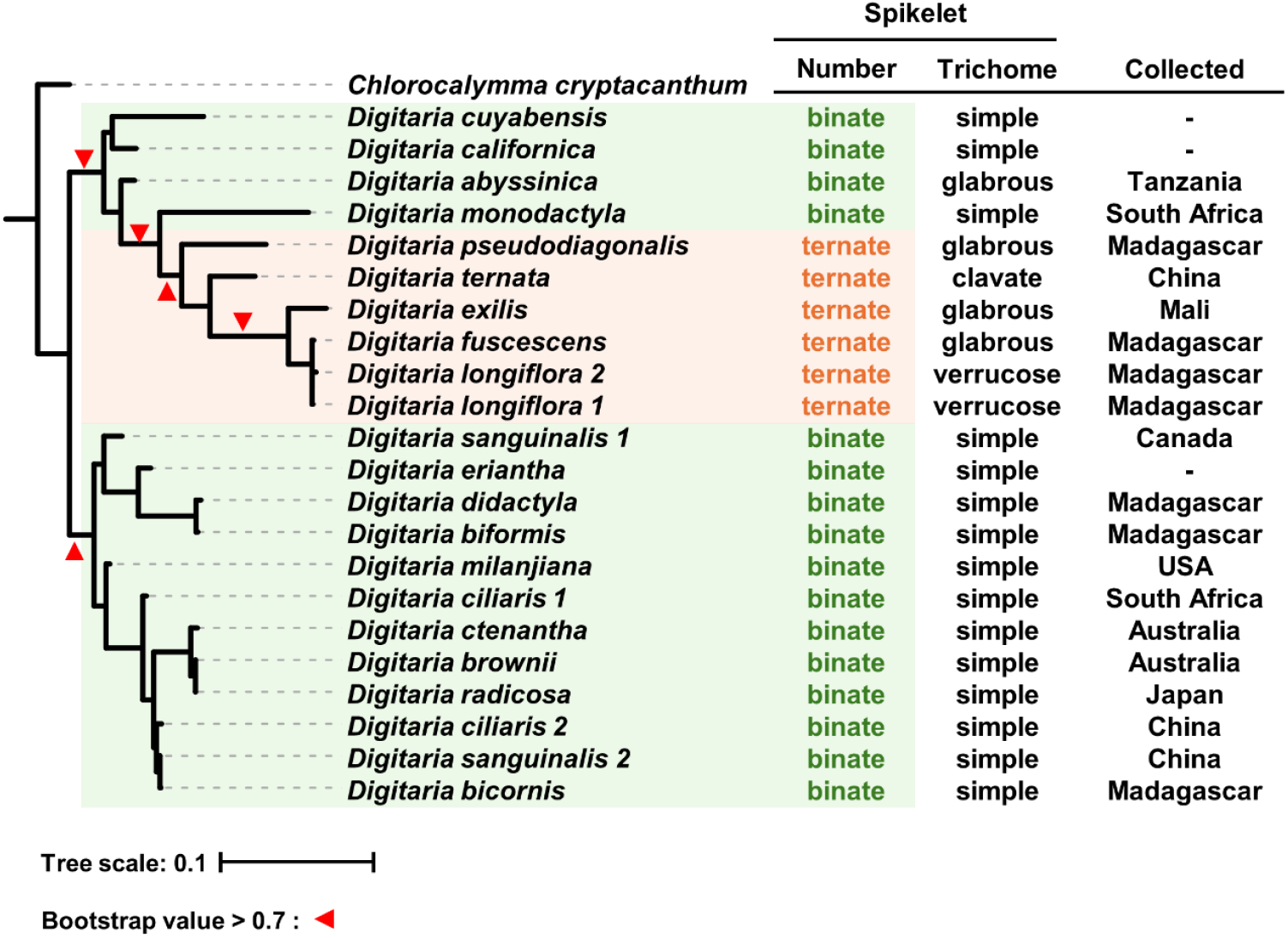
The phylogenetic relationships of *Digitaria* species reflect their spikelet morphology. Phylogenetic tree of 19 *Digitaria* species and one *Chlorocalymma* species as an outgroup. The maximum-likelihood phylogenetic tree was generated in RAxML version 8.2.12 with the JTT model using 2,052 SNPs. The scale bar indicates the evolutionary distance in nucleotide substitution per site. The well-supported nodes are visualized as red arrowheads with bootstrap values > 0.7 based on 1000 iterations. The number of spikelets at each node on the rachis is indicated by different background colors. The right table indicates the number of spikelets at each node on the rachis, the trichome morphology, and country where each sample was collected.

Our phylogenetic tree included multiple samples for three *Digitaria* species. The two *D. longiflora* plants collected from Madagascar had almost identical genotypes (**Figure 1**). However, the two *D. sanguinalis* plants collected from Canada or China and the *D. ciliaris* plants collected from China or South Africa were phylogenetically more diverse but still fell within the same clade (**Figure 1**).

To identify the relationship between genetic variation and phenotypic variation, we investigated spikelet morphology, which is traditionally used for the grouping of *Digitaria* species (Henrard 1950). The number of spikelets at each node on the rachis was clearly different among the two clades defined by the phylogenetic tree (**Figure 1**). Five *Digitaria* species (*D. pseudodiagonalis, D. ternata, D. exilis, D. fuscescens*, and *D. longiflora*) constituted a well-supported clade with ternate spikelets (**Figure 1**). Other species showed binate spikelets (**Figure 1**). Looking at trichome morphology of spikelets, *Digitaria* species from the second clade (*D. sanguinalis* to *D. bicornis*) had simple hair form. By contrast, *Digitaria* species from the first clade (*D. cuyabensis* to *D. longiflora*) showed wide variation in their spikelet trichome morphology (**Figure 1**). These results indicate that a phylogenetic analysis based on the *D. radicosa* genome may help to elucidate the genetic relatedness among *Digitaria* species and among plants of the same species, as well as the evolution of spikelet morphology.

### Genetic diversity of *Digitaria* species in Japan

Considering the wide geographical distribution of *Digitaria* species in Japan and the reported phenotypic variation among *D. ciliaris* plants, we hypothesized that this species may comprise genetically distinct subpopulations (Kataoka *et al*. 1986; Tsuyuzaki 2005; Fukano *et al*. 2020). To investigate the genetic differentiation among geographically distinct areas and habitats, we performed a principal component analysis (PCA) using SNPs obtained from 10 *D. radicosa* plants and 11 *D. ciliaris* plants collected from the Kansai region, Japan (**Figure 2a**). We obtained *D. radicosa* samples from all four locations to reveal any underlying population structure associated with geographical distribution. We also collected *D. ciliaris* samples from three habitats (parks, paddy fields, and coastal areas) to investigate genetic variation among plants growing in different habitats (**Figure 2c, Figure S1**). The PCA showed that two plants were likely misidentified as *D. ciliaris*, which we removed for the downstream analysis (**Figure S2**). A total of 206 and 245 SNPs were identified from among plant samples belonging to *D. radicosa* (10 samples) and *D. ciliaris* (9 samples), respectively, and 101 SNPs were identified from among all samples from both species.

**Figure 2.**
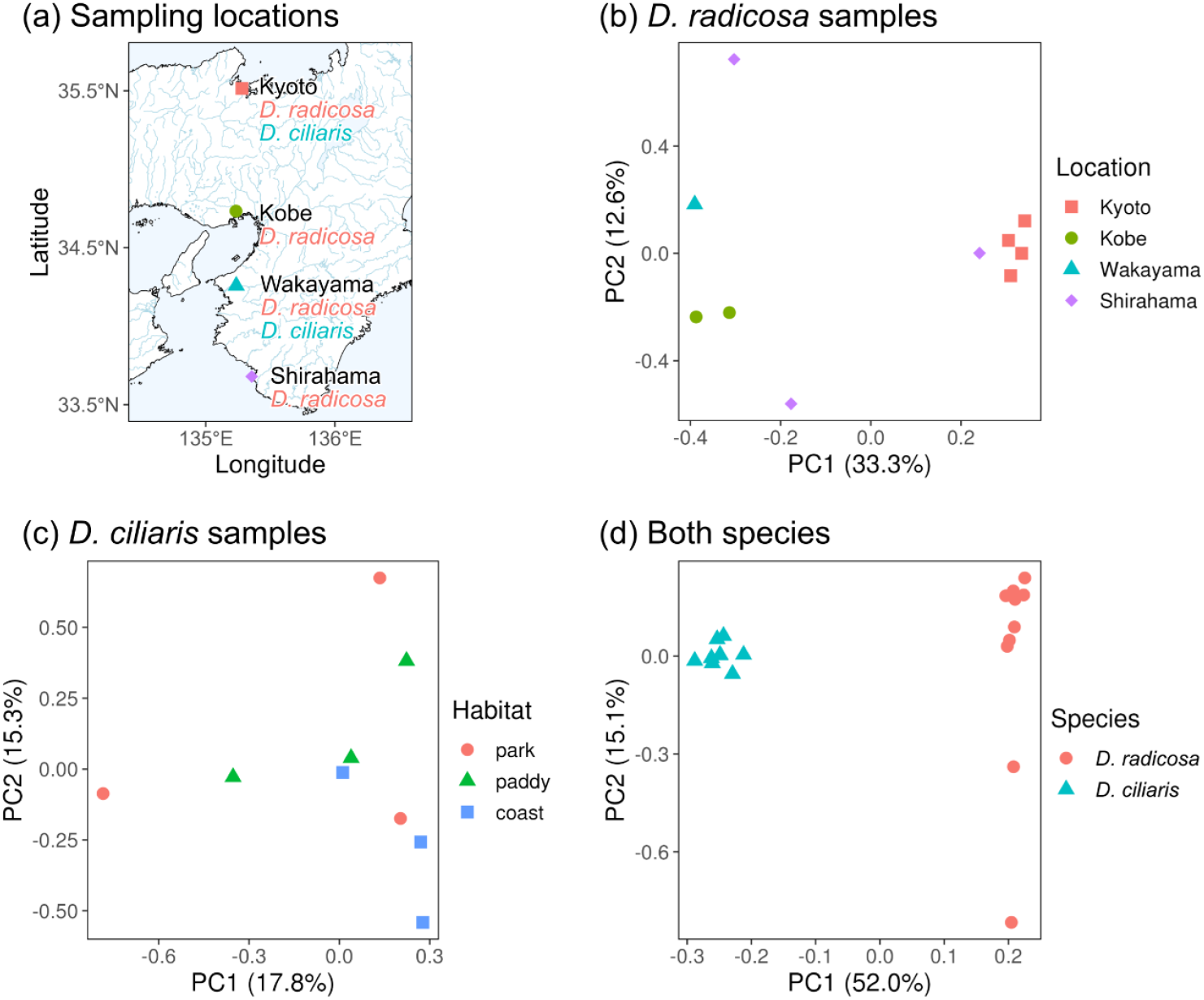
Population genetic analysis of *Digitaria radicosa* and *D. ciliaris*. (a) Sampling locations of *D. radicosa* and *D. ciliaris* plants. The map is based on Global Map Japan (Geospatial Information Authority of Japan 2016). (b– d) Principal component analysis (PCA) for *D. radicosa* plants sampled from four locations (b), *D. ciliaris* plants sampled separately from different habitats at each sampling location (c), and plants from both species (d). The numbers in parentheses indicate the percentage of standing variance explained by each principal component (PC).

The *D. radicosa* plants formed two clusters along principal component 1 (PC1), which explained 33.3% of the standing genomic variation (**Figure 2b**). Plants collected from the same location clustered together (Kyoto and Kobe) or were far from each other (Shirahama), indicating little genetic variation among plants from Kyoto or Kobe and large genetic variation among plants originating from Shirahama.

*D. ciliaris* plants collected from different habitats showed a loose clustering pattern (**Figure 2c**). Plants from different habitats partially overlapped in the PCA plot, although plants collected from paddy fields and coastal areas appeared to be separated along PC2. Plants collected from parks showed the largest variation along PC1 and PC2.

A PCA plot using all *D. radicosa* and *D. ciliaris* plants supported the clear genomic separation of these two species (**Figure 2d**). Indeed, the two species formed distinct clusters along PC1, explaining 52.0% of the total genomic variation. While all *D. ciliaris* plants were contained within a single cluster, two *D. radicosa* plants were distant from the other eight plants of the same species along PC2 (**Figure 2d**).

### Comparative analysis

Since the only available chromosome-scale genome assembly is for the cultivated *Digitaria* species *D. exilis*, we did not investigate genomic structural variation between wild and cultivated *Digitaria* species (Abrouk *et al*. 2020; Wang *et al*. 2021). To explore the presence of structural variation and the degree of synteny between wild *Digitaria* and cultivated *Digitaria* species, we performed a synteny analysis between *D. radicosa* and *D. exilis* using their respective high-quality, chromosome-scale genome assemblies. We also identified syntenic blocks between *D. radicosa* and rice to investigate conserved genomic regions between the two Poaceae species. Synteny analysis revealed a high degree of synteny at the whole chromosome level between *D. radicosa* and *D. exilis* across all chromosomes, aside from chromosomal translocations observed at the tip of chromosome 2 and the end of chromosome 7 in *D. radicosa* (**Figure 3a**). A similar analysis between the *D. radicosa* and rice genomes revealed conserved syntenic blocks as chromosome scale among chromosomes 1, 4, 6, and 8 of *D. radicosa* and chromosomes 2, 6, 8, and 11 of rice, respectively (**Figure 3b**). Other *D. radicosa* chromosomes also showed sequence conservation with some genomic regions in the rice genome, although these conserved sequences were spread across several chromosomes (**Figure 3b**). These results indicate that the *D. radicosa* genome may have genes orthologous to functionally important genes of Poaceae species including fonio and rice.

**Figure 3.**
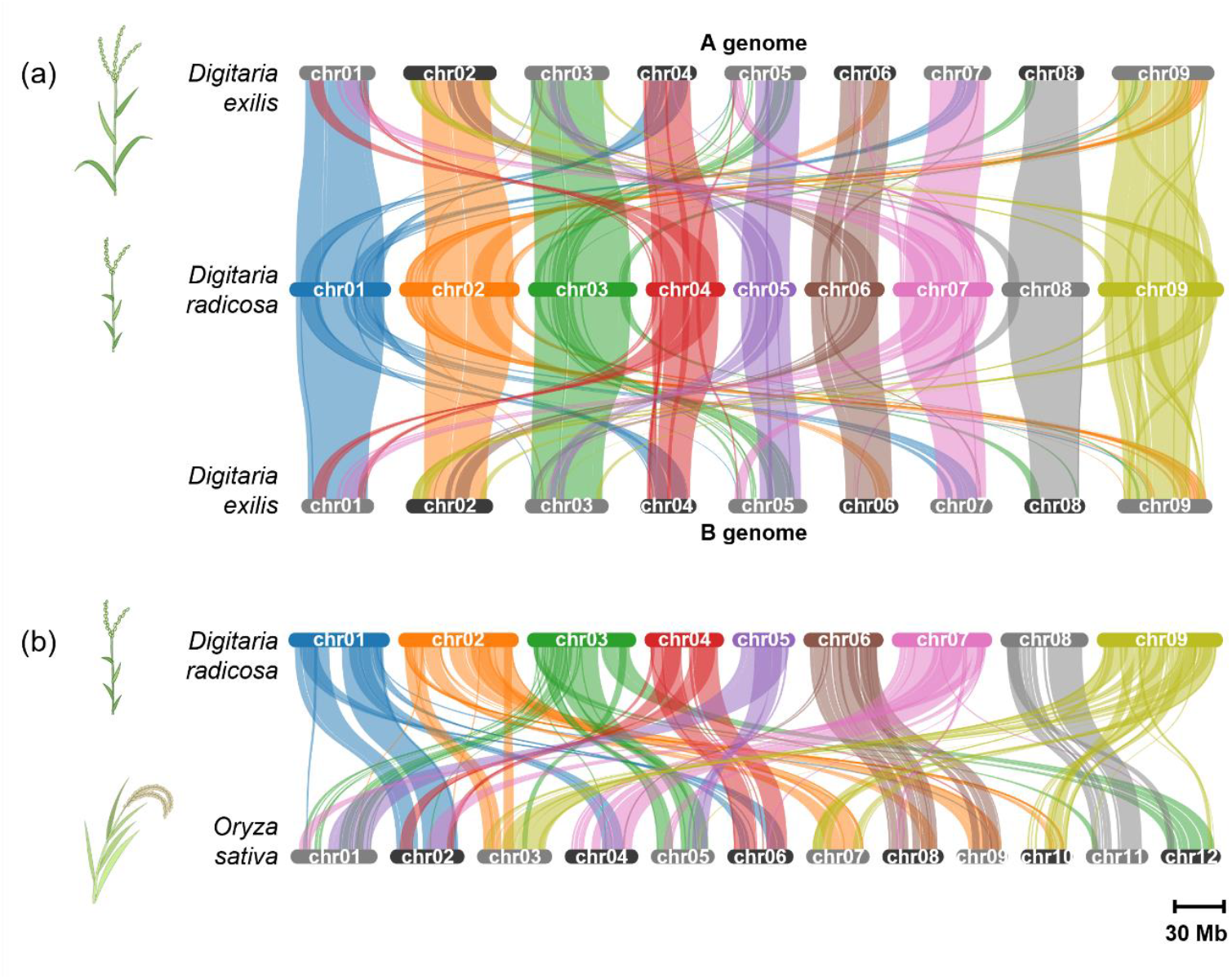
The *Digitaria radicosa* reference genome is highly syntenic with the *Digitaria exilis* and rice genomes. (a) The top plot shows the syntenic blocks between *D. radicosa* and the A or B genome of *D. exilis*. (b) The bottom plot shows the syntenic blocks between *D. radicosa* and Nipponbare cultivar of rice. Syntenic blocks were identified by MCScanX and visualized by SynVisio.

### Resistance genes in the *Digitaria radicosa* genome

NLR proteins recognize effectors secreted by blast fungus and induce an immune response in Poaceae species including rice (Jones *et al*. 2016). Previous studies have shown how domains within NLRs, named integrated domains (IDs), are involved in effector recognition (Kroj *et al*. 2016; Marchal *et al*. 2022). We hypothesized that *Digitaria* species might have NLRs with IDs recognizing blast fungus isolates from rice plants. To investigate NLRs in *D. radicosa*, we extracted 155 genes annotated as NLR genes from our reference genome and performed a phylogenetic analysis on their predicted protein sequences to classify them, together with 100 functionally validated NLR proteins from 14 other Poaceae species. We determined that *D. radicosa* NLRs are phylogenetically diverse (**Figure S3**). Combined with the results of synteny analysis, we identified *D. radicosa* genes orthologous to the rice NLR genes *RESISTANCE GENE ANALOG 5* (*RGA5*) and *RGA4*, as well as orthologs for *PYRICULARIA ORYZAE RESISTANCE K-2* (*Pik-2*) on chromosome 8 of *D. radicosa* (**Figure 4**) (Kanzaki *et al*. 2012; Cesari *et al*. 2013). These rice NLRs recognize effectors secreted by *M. oryzae* and induce immune responses in rice. In addition, one putative *RGA5* ortholog in *D. radicosa* harbored an Exo70 domain in its C terminus as an ID (**Figure 4**). These results suggest that the *D. radicosa* genome encodes NLRs that may recognize *M. oryzae* rice isolates, making the *Digitaria* genome a potentially useful resource for identifying resistance genes to blast fungus that can be deployed in other Poaceae crops.

**Figure 4.**
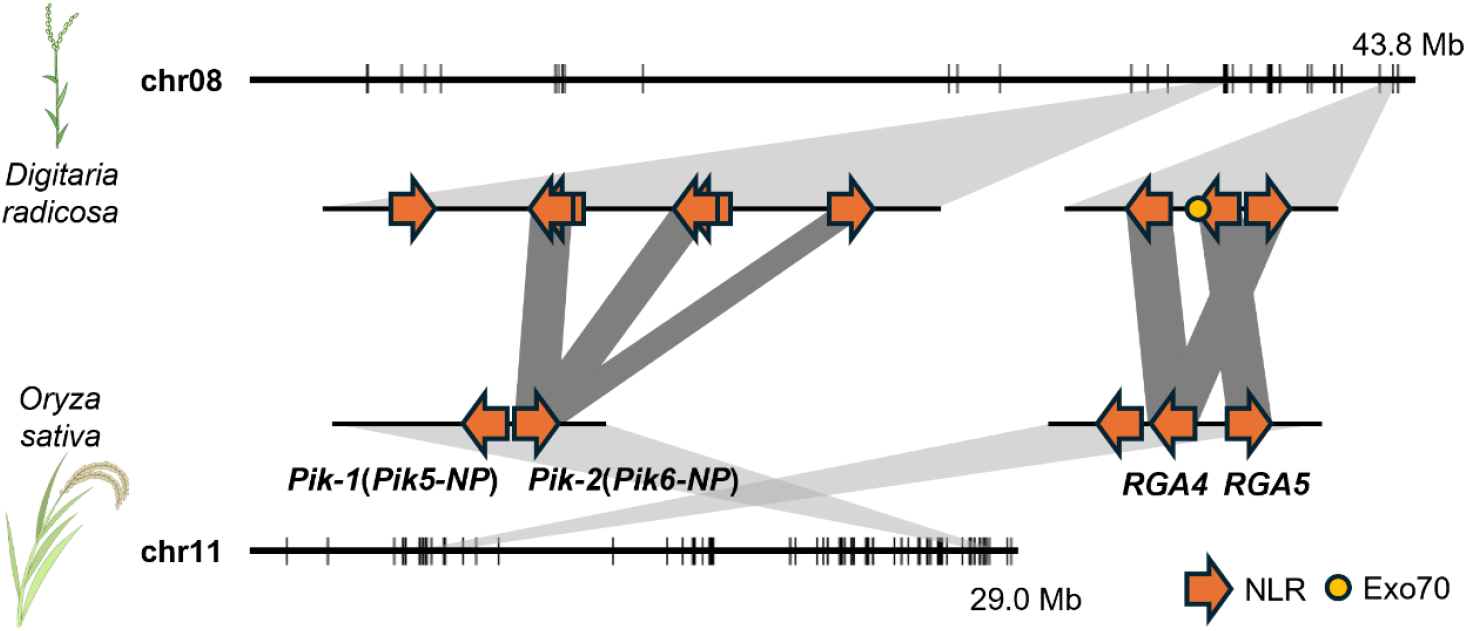
Examples of functionally validated rice NLR orthologs in *Digitaria radicosa*. Top, chromosome 8 of *D. radicosa*; bottom, chromosome 11 of rice. Annotated NLRs are displayed as vertical lines. Middle, genomic regions of *D. radicosa* carrying NLR genes orthologous to *Pik-2* (*Pik6-NP* in Nipponbare) and *RGA4/5* from rice. Orange arrows indicate NLRs; the light orange circle indicates an integrated domain. Dark gray shaded region between NLRs indicates phylogenetically well-supported orthologs.

## Discussion

We generated a chromosome-scale reference genome assembly of the diploid species *D. radicosa*. We obtained nine pseudochromosomes with N50 scaffold values and BUSCO scores supporting the high contiguity and integrity of this new genome assembly (**Table 1**). The only previously available chromosome-scale reference genome was from a tetraploid *Digitaria* species, *D. exilis*, with no reference genome for a diploid *Digitaria* species (Abrouk *et al*. 2020; Wang *et al*. 2021). Most standard tools for genomic analysis have been developed for diploid genomes and are not easily applied to polyploid genomes (Dufresne *et al*. 2014). Therefore, our assembled diploid *Digitaria* genome will allow population genetic analysis and evolutionary analysis of *Digitaria* species.

To verify the utility of our newly assembled diploid *Digitaria* genome, we elucidated the genetic relatedness of 19 *Digitaria* species, resolving their previously inconsistent classification (**Figure 1**). For instance, the species *D. monodactyla* with binate spikelets was previously classified together with species having ternate spikelets (Touafchia *et al*. 2023). However, our phylogenetic tree based on genome-wide SNPs demonstrated that *Digitaria* species with ternate spikelet morphology constitute one well-supported clade, suggesting a clear genetic difference between species with ternate and binate spikelet morphology (**Figure 1**). Therefore, our evolutionary analysis based on diploid *Digitaria* species possibly provides a more precise classification of *Digitaria* species. In addition, our phylogenetic tree showed distant genetic relatedness among some plants from the same species collected from different geographical locations (**Figure 1**). This suggests that geographical isolation exerts a significant effect on the genetic diversity of *Digitaria* species. Alternatively, this might have resulted from incorrect genotyping for several polyploid species that require high coverage to genotype correctly (Gerard *et al*. 2018; Gould 1963; Kaur and Gupta 2016; Koul and Gohil 1991). Published sequencing data for eight *Digitaria* species were generated using target enrichment (Baker *et al*. 2021). These sequencing data were therefore restricted in their genome coverage and do not offer a comprehensive view of genomic diversity among *Digitaria* species. Phylogenetic analysis based on sequencing data for the entire genome of *Digitaria* species might provide more precise genetic classification of *Digitaria* species (Leaché and Oaks 2017).

We investigated the genetic structure of *Digitaria* species in Japan, among plants from the same species (intraspecific) and among different species (interspecific). As all *D. radicosa* plants were separated into two clusters by PCA, we speculate that the samples we collected are possibly composed of two genetically distinct subpopulations (**Figure 2b**). The genetic difference based on 206 SNPs between these two clusters might not be associated with their geographical distribution, since plants collected from Shirahama were present into both clusters (**Figure 2b**). In addition, plants collected from Kyoto and Kobe each formed a tight cluster, suggesting that these samples might be inbred or of clonal origin (**Figure 2b**). Therefore, in Kyoto and Kobe, we might have sampled only one subpopulation, which does not preclude the existence of diverse subpopulations not yet sampled.

*D. ciliaris* differentiates into divergent populations adapted to different environmental habitats (Kataoka *et al*. 1986; Tsuyuzaki 2005; Fukano *et al*. 2020). However, our results showed a loose clustering of *D. ciliaris* plant samples, suggesting the absence of clear population structure as a function of the environment (**Figure 2c**). Therefore, in addition to parallel evolution of adaptive traits in each local population, frequent gene flow among different habitats might have also occurred in *D. ciliaris*. Gene flow fostered by the outcrossing capability of *D. ciliaris* may have maintained genetic variation of each locally adapted population under adaptive selection, as hypothesized for herbicide-resistant populations of several weed species, including *D. insularis* (Kataoka and Kataoka 1991; Karn and Jasieniuk 2017; Gonçalves Netto *et al*. 2021; Brunharo and Tranel 2023).

In contrast to the unclear intraspecific genetic structure, we observed a clear species boundary between *D. radicosa* and *D. ciliaris*. As clusters comprising these two species were clearly distinguished with no intermediate samples, there is little possibility of interspecific hybridization (**Figure 2d**). Difference in ploidy level may have worked as a reproductive barrier to keep these two species isolated (Hirayoshi and Yasue 1955).

We conclude that population genetic analysis based on our newly assembled reference genome succeeded in identifying two distinct subpopulations in *D. radicosa* and highlighted the different interspecific genetic structure of *Digitaria* species, but a more detailed analysis with a larger sample size will be required to unveil the population structure of *Digitaria* species across Japan.

Synteny analysis between *D. radicosa* and *D. exilis* identified a high degree of syntenic blocks across entire chromosomes, except for two genome rearrangements (**Figure 3**). Our evolutionary analysis showed that *D. radicosa* and *D. exilis* belong to different well-supported clades (**Figure 1**). Therefore, these genome rearrangements might have occurred after the clades diverged. In addition, synteny analysis revealed that most rice genomic regions have syntenic blocks with some regions of the *D. radicosa* genome (**Figure 3**). Since the genus *Digitaria* and the genus *Oryza* diverged almost 50 million years ago, this synteny may indicate that most core gene sets from Poaceae species are conserved in *D. radicosa* and rice (Pessoa-Filho *et al*. 2017). Comparative genomics including synteny analysis can help identify putative functions of genes (Bennetzen and Chen 2008; Wijerathna-Yapa *et al*. 2023). Therefore, the gene functions of Poaceae crops may be elucidated from these highly conserved synteny blocks between *D. radicosa* and Poaceae crops.

We identified NLR genes in the *D. radicosa* genome as representative resistance genes. *D. radicosa* has genes orthologous to rice *RGA4/5*, which recognize effectors secreted by *M. oryzae* and induce an immune response (**Figure 4**) (Kanzaki *et al*. 2012; Cesari *et al*. 2013). Interestingly, the *RGA5* ortholog in *D. radicosa* carries the ID Exo70 in its C terminus (**Figure 4**). *RGA5* orthologs in *Oryza* species contain a wide variety of IDs and are thought to recognize corresponding effectors (Shimizu *et al*. 2022). In addition, Exo70 interacts with the *Pii-1/2* NLR pair to recognize the *M. oryzae* Avr-Pii effector to induce immune responses (Takagi *et al*. 2013; Fujisaki *et al*. 2017). Therefore, *RGA5* orthologs from *D. radicosa* may recognize effectors secreted by *M. oryzae* through their Exo70 domain. *D. radicosa* also has genes orthologous to a wide variety of functionally validated NLRs in Poaceae species (**Figure S3**). These results suggest that *D. radicosa* could be a useful genomic resource for improving plant immunity in Poaceae crops.

In summary, we constructed a chromosome-scale reference genome for the diploid species *D. radicosa*. This newly assembled diploid *Digitaria* genome allowed us to perform standard genomic analyses of *Digitaria* species. Our evolutionary analysis of 19 *Digitaria* species based on our new reference genome illustrated their extent of genetic relatedness. Population genetic analysis of *D. radicosa* and *D. ciliaris* revealed the genetic structure within species and the relationships between them. Larger-scale population genetic analysis will provide more valuable insights into the genetic relatedness of *Digitaria* species. Furthermore, the *D. radicosa* genome showed highly conserved syntenic blocks with the representative Poaceae crop and harbored putative resistance genes for *M. oryzae*. These findings indicate that our new diploid *Digitaria* genome could be a useful resource for genomic studies of *Digitaria* species and for Poaceae crop breeding.

## Data availability

PacBio HiFi sequencing reads and short reads generated for the reference genome have been deposited in the SRA of NCBI under BioProject PRJNA1097755. The assembled genome of *D. radicosa* was also deposited at NCBI under BioProject PRJNA1097755. MIG-seq sequencing data generated for population genetic analysis have been deposited in the SRA under BioProject PRJNA1098455. The sequences of the reference genome and gene annotation data were deposited in Zenodo (10.5281/zenodo.10952520). All scripts and commands used in this study were deposited in a GitHub repository (https://github.com/slt666666/D_radicosa_genome).

## Supplemental Data

Figure S1. Example of Digitaria radicosa plant and examples of Digitaria ciliaris plants growing in three habitats.

Figure S2. Principal component analysis of all samples before removing putatively misidentified samples.

Figure S3. Phylogenetic tree of identified NLR proteins from Digitaria radicosa and functionally validated NLRs from Poaceae species.

Table S1. Sampling information of *Digitaria radicosa* and *Digitaria ciliaris*.

Table S2. Accession numbers of sequencing data for gene annotation and evolutionary analysis.

Table S3. Summary statistics of repeat annotation.

Table S4. Summary statistics of OMArk analysis.

File S1. Genetic variants data and phylogenetic tree file used for Figure 1.

File S2. Genetic variants data used for Figure 2 and S2.

File S3. Amino acid sequence, alignment, and phylogenetic tree files used for Figure 4 and S3.

## Acknowledgements

Computations were partially performed on the NIG supercomputer at RIOS National Institute of Genetics. We thank Hiroshi Tsuyuzaki (Faculty of Bioresource Sciences, Akita Prefectural University, Japan) for the assistance in species identification. We thank Luna Yamamori (Seto Marine Biological Laboratory, Kyoto University, Japan) for illustrations on figures.

## Literature Cited

Abrouk M, Ahmed HI, Cubry P, Šimoníková D, Cauet S, et al. 2020. Fonio millet genome unlocks African orphan crop diversity for agriculture in a changing climate. Nat. Commun. 11: 4488.

Alonge M, Lebeigle L, Kirsche M, Jenike K, Ou S, et al. 2022. Automated assembly scaffolding using RagTag elevates a new tomato system for high-throughput genome editing. Genome Biol. 23: 258.

Ayenan MAT, Sodedji KAF, Nwankwo CI, Olodo KF, Alladassi MEB. 2018 Harnessing genetic resources and progress in plant genomics for fonio (Digitaria spp.) improvement. Genet. Resour. Crop Evol. 65: 373–386.

Baker WJ, Dodsworth S, Forest F, Graham SW, Johnson MG, et al. 2021. Exploring Angiosperms353: An open, community toolkit for collaborative phylogenomic research on flowering plants. Am. J. Bot. 108: 1059–1065.

Bandi V, Gutwin C, Siri JN, Neufeld E, Sharpe A, et al. 2022. Visualization Tools for Genomic Conservation, pp. 285–308 in Plant Bioinformatics: Methods and Protocols, edited by D. Edwards. Springer US, New York, NY.

Bennetzen JL, Chen M. 2008 Grass Genomic Synteny Illuminates Plant Genome Function and Evolution. Rice 1: 109–118.

Brunharo CACG, Tranel PJ. 2023 Repeated evolution of herbicide resistance in Lolium multiflorum revealed by haplotype-resolved analysis of acetyl-CoA carboxylase. Evol. Appl. 16: 1969–1981.

Cesari S, Thilliez G, Ribot C, Chalvon V, Michel C, et al. 2013. The rice resistance protein pair RGA4/RGA5 recognizes the Magnaporthe oryzae effectors AVR-Pia and AVR1-CO39 by direct binding. Plant Cell 25: 1463–1481.

Chang CC, Chow CC, Tellier LC, Vattikuti S, Purcell SM, et al. 2015. Second-generation PLINK: rising to the challenge of larger and richer datasets. Gigascience 4: 7.

Chen S, Zhou Y, Chen Y, Gu J. 2018 fastp: an ultra-fast all-in-one FASTQ preprocessor. Bioinformatics 34: i884–i890.

Cheng H, Concepcion GT, Feng X, Zhang H, Li H. 2021 Haplotype-resolved de novo assembly using phased assembly graphs with hifiasm. Nat. Methods 18: 170–175.

Danecek P, Auton A, Abecasis G, Albers CA, Banks E, et al. 2011. The variant call format and VCFtools. Bioinformatics 27: 2156–2158.

Darriba D, Posada D, Kozlov AM, Stamatakis A, Morel B, et al. 2020. ModelTest-NG: A New and Scalable Tool for the Selection of DNA and Protein Evolutionary Models. Mol. Biol. Evol. 37: 291–294.

Dobin A, Davis CA, Schlesinger F, Drenkow J, Zaleski C, et al. 2013. STAR: ultrafast universal RNA-seq aligner. Bioinformatics 29: 15–21.

Dufresne F, Stift M, Vergilino R, Mable BK. 2014 Recent progress and challenges in population genetics of polyploid organisms: an overview of current state-of-the-art molecular and statistical tools. Mol. Ecol. 23: 40–69.

Flynn JM, Hubley R, Goubert C, Rosen J, Clark AG, et al. 2020. RepeatModeler2 for automated genomic discovery of transposable element families. Proc. Natl. Acad. Sci. U. S. A. 117: 9451–9457.

Fujisaki K, Abe Y, Kanzaki E, Ito K, Utsushi H, et al. 2017. An unconventional NOI/RIN4 domain of a rice NLR protein binds host EXO70 protein to confer fungal immunity. bioRxiv 239400.

Fukano Y, Guo W, Uchida K, Tachiki Y. 2020 Contemporary adaptive divergence of plant competitive traits in urban and rural populations and its implication for weed management. J. Ecol. 108: 2521–2530.

Gabriel L, Brůna T, Hoff KJ, Ebel M, Lomsadze A, et al. 2023. BRAKER3: Fully automated genome annotation using RNA-Seq and protein evidence with GeneMark-ETP, AUGUSTUS and TSEBRA. bioRxiv 2023.06.10.544449.

Gerard D, Ferrão LFV, Garcia AAF, Stephens M. 2018 Genotyping Polyploids from Messy Sequencing Data. Genetics. 210: 789–807.

Global Map Japan. 2016 Tokyo: Geospatial Information Authority of Japan; [accessed 2024 Mar 24]. https://www.gsi.go.jp/kankyochiri/gm_jpn.html

Gonçalves Netto A, Cordeiro EM, Nicolai M, de Carvalho SJ, Ovejero RFL, et al. 2021. Population genomics of Digitaria insularis from soybean areas in Brazil. Pest Manag. Sci. 77: 5375–5381.

Gould FW. 1963 Cytotaxonomy of Digitaria sanguinalis and D. adscendens. Brittonia. 15:241–244.

Guan D, McCarthy SA, Wood J, Howe K, Wang Y, et al. 2020. Identifying and removing haplotypic duplication in primary genome assemblies. Bioinformatics 36: 2896–2898.

Hirayoshi I, Yasue T. 1955 Cytogenetical Studies on forage plants. (II) : Chromosome numbers and some Specific characters in Japanese native species of Digitaria. Japanese Journal of Breeding 5: 47–48.

Hu J, Wang Z, Liang F, Liu SL, Ye K, et al. 2024. NextPolish2: A Repeat-aware Polishing Tool for Genomes Assembled Using HiFi Long Reads. Genomics Proteomics Bioinformatics.

Jideani IA. 1999 Traditional and possible technological uses of Digitaria exilis (acha) and Digitaria iburua (iburu): a review. Plant Foods Hum. Nutr. 54: 363–374.

Jones JDG, Vance RE, Dangl JL. 2016 Intracellular innate immune surveillance devices in plants and animals. Science 354:.

Kanzaki H, Yoshida K, Saitoh H, Fujisaki K, Hirabuchi A, et al. 2012. Arms race co-evolution of Magnaporthe oryzae AVR-Pik and rice Pik genes driven by their physical interactions. Plant J. 72: 894–907.

Karn E, Jasieniuk M. 2017 Genetic diversity and structure of Lolium perenne ssp. Multiflorum in California vineyards and orchards indicate potential for spread of herbicide resistance via gene flow. Evol. Appl. 10: 616–629.

Kataoka M, Ibaraki K, Tokunaga H. 1986 Differential Heading Behavior of Some Digitaria adscendens Henr. Populations. Weed Technol. 31: 36–40.

Kataoka M, Kataoka K. 1991 Lower-Lemma Seta Inheritance and Outcrossing Rate of Digitaria adscendens. Weed Technol. 36: 82–84.

Katoh K, Standley DM. 2013 MAFFT multiple sequence alignment software version 7: improvements in performance and usability. Mol. Biol. Evol. 30: 772–780.

Kaur N, Gupta RC. 2016 Cytological Study in Some Members of Tribe Paniceae (Poaceae) from Rajasthan. Cytologia. 81:13–17.

Koul KK, Gohil RN. 1991 Cytogenetic Studies on Some Kashmir Grasses. VIII. Tribe Agrostideae, Festuceae and Paniceae. Cytologia. 56:437–452.

Kourelis J, Sakai T, Adachi H, Kamoun S. 2021 RefPlantNLR is a comprehensive collection of experimentally validated plant disease resistance proteins from the NLR family. PLoS Biol. 19: e3001124.

Kroj T, Chanclud E, Michel-Romiti C, Grand X, Morel JB. 2016 Integration of decoy domains derived from protein targets of pathogen effectors into plant immune receptors is widespread. New Phytol. 210: 618–626.

Leaché AD, Oaks JR. 2017 The utility of single nucleotide polymorphism (SNP) data in phylogenetics. Annu. Rev. Ecol. Evol. Syst. 48: 69–84.

Li H. 2011 A statistical framework for SNP calling, mutation discovery, association mapping and population genetical parameter estimation from sequencing data. Bioinformatics 27: 2987–2993.

Li H. 2013 Aligning sequence reads, clone sequences and assembly contigs with BWA-MEM. arXiv [q-bio.GN].

Li H, Handsaker B, Wysoker A, Fennell T, Ruan J, et al. 2009. The Sequence Alignment/Map format and SAMtools. Bioinformatics 25: 2078–2079.

Manni M, Berkeley MR, Seppey M, Simão FA, Zdobnov EM. 2021 BUSCO Update: Novel and Streamlined Workflows along with Broader and Deeper Phylogenetic Coverage for Scoring of Eukaryotic, Prokaryotic, and Viral Genomes. Mol. Biol. Evol. 38: 4647–4654

Maqbool A, Saitoh H, Franceschetti M, Stevenson CEM, Uemura A, et al. 2015. Structural basis of pathogen recognition by an integrated HMA domain in a plant NLR immune receptor. Elife 4:.

Marchal C, Michalopoulou VA, Zou Z, Cevik V, Sarris PF. 2022 Show me your ID: NLR immune receptors with integrated domains in plants. Essays Biochem. 66: 527–539.

McKenna A, Hanna M, Banks E, Sivachenko A, Cibulskis K, et al. 2010. The Genome Analysis Toolkit: a MapReduce framework for analyzing next-generation DNA sequencing data. Genome Res. 20: 1297–1303.

Nevers Y, Warwick Vesztrocy A, Rossier V, Train CM, Altenhoff A, et al. 2024. Quality assessment of gene repertoire annotations with OMArk. Nat. Biotechnol.

Ou SH. 1980 Pathogen Variability and Host Resistance in Rice Blast Disease. Annu. Rev. Phytopathol. 18: 167–187.

Pessoa-Filho M, Martins AM, Ferreira ME. 2017 Molecular dating of phylogenetic divergence between Urochloa species based on complete chloroplast genomes. BMC Genomics 18: 516.

Pitman WD, Chambliss CG, Hacker JB. 2016 Digitgrass and other species of Digitaria, pp. 715–743 in Warm-Season (C4) Grasses, Agronomy Monographs, American Society of Agronomy, Crop Science Society of America, Soil Science Society of America, Madison, WI, USA.

Prance SG, Nesbitt M. 2004 The Cultural History of Plants. Routledge.

Ravet K, Patterson EL, Krähmer H, Hamouzová K, Fan L, et al. 2018. The power and potential of genomics in weed biology and management. Pest Manag. Sci. 74: 2216–2225.

Sakai H, Lee SS, Tanaka T, Numa H, Kim J, et al. 2013. Rice Annotation Project Database (RAP-DB): an integrative and interactive database for rice genomics. Plant Cell Physiol. 54: e6.

Scarcelli N, Barnaud A, Eiserhardt W, Treier UA, Seveno M, et al. 2011. A set of 100 chloroplast DNA primer pairs to study population genetics and phylogeny in monocotyledons. PLoS One 6: e19954.

Shimizu M, Hirabuchi A, Sugihara Y, Abe A, Takeda T, et al. 2022. A genetically linked pair of NLR immune receptors shows contrasting patterns of evolution. Proc. Natl. Acad. Sci. U. S. A. 119: e2116896119.

Soreng RJ, Peterson PM, Zuloaga FO, Romaschenko K, Clark LG, et al. 2022. A worldwide phylogenetic classification of the Poaceae (Gramineae) III: An update. J. Syst. Evol. 60: 476–521.

Stamatakis A. 2006 RAxML-VI-HPC: maximum likelihood-based phylogenetic analyses with thousands of taxa and mixed models. Bioinformatics 22: 2688–2690.

Suyama Y, Matsuki Y. 2015 MIG-seq: an effective PCR-based method for genome-wide single-nucleotide polymorphism genotyping using the next-generation sequencing platform. Sci. Rep. 5: 16963.

Takagi H, Uemura A, Yaegashi H, Tamiru M, Abe A, et al. 2013. MutMap-Gap: whole-genome resequencing of mutant F2 progeny bulk combined with de novo assembly of gap regions identifies the rice blast resistance gene Pii. New Phytol. 200: 276–283.

Tarailo-Graovac M, Chen N. 2009 Using RepeatMasker to identify repetitive elements in genomic sequences. Curr. Protoc. Bioinformatics Chapter 4: 4.10.1–4.10.14.

Tosa Y, Chuma I. 2014 Classification and parasitic specialization of blast fungi. J. Gen. Plant Pathol. 80: 202–209.

Touafchia S, Maurin O, Boonsuk B, Hodkinson TR, Chantaranothai P, et al. 2023. Evolutionary history, traits, and weediness in Digitaria (Poaceae: Panicoideae). Bot. J. Linn. Soc. 203: 1–19.

Tsuyuzaki H. 2005 Adaptation of morphological and ecological characteristics of Digitaria ciliaris (Retz.) Koeler in adjacent habitats. Journal of Weed Science and Technology 50: 10–17.

Vasimuddin M, Misra S, Li H, Aluru S. 2019 Efficient Architecture-Aware Acceleration of BWA-MEM for Multicore Systems, pp. 314–324 in 2019 IEEE International Parallel and Distributed Processing Symposium (IPDPS), IEEE.

Vega AS, Rua GH, Fabbri LT, Rúgolo de Agrasar ZE. 2009 A Morphology-Based Cladistic Analysis of Digitaria (Poaceae, Panicoideae, Paniceae). Syst. Bot. 34: 312–323.

Wang X, Chen S, Ma X, Yssel AEJ, Chaluvadi SR, et al. 2021. Genome sequence and genetic diversity analysis of an under-domesticated orphan crop, white fonio (Digitaria exilis). Gigascience 10:.

Wang Y, Tang H, Debarry JD, Tan X, Li J, et al. 2012. MCScanX: a toolkit for detection and evolutionary analysis of gene synteny and collinearity. Nucleic Acids Res. 40: e49.

Weinert-Nelson JR, Meyer WA, Williams CA. 2022 Crabgrass as an equine pasture forage: impact of establishment method on yield, nutrient composition, and horse preference. Transl Anim Sci 6: txac050.

Wijerathna-Yapa A, Bishnoi R, Ranawaka B, Maya Magar M, Ur Rehman H, et al. 2023. Rice– wheat comparative genomics: Gains and gaps. The Crop Journal.

Yoshida K, Saunders DGO, Mitsuoka C, Natsume S, Kosugi S, et al. 2016 Host specialization of the blast fungus Magnaporthe oryzae is associated with dynamic gain and loss of genes linked to transposable elements. BMC Genomics 17: 370.

Zimin AV, Marçais G, Puiu D, Roberts M, Salzberg SL, et al. 2013. The MaSuRCA genome assembler. Bioinformatics 29: 2669–2677.

