## Supplemental Figures for "A chromosome-scale genome assembly of Timorese crabgrass (*Digitaria radicosa*): a useful genomic resource for the Poaceae"

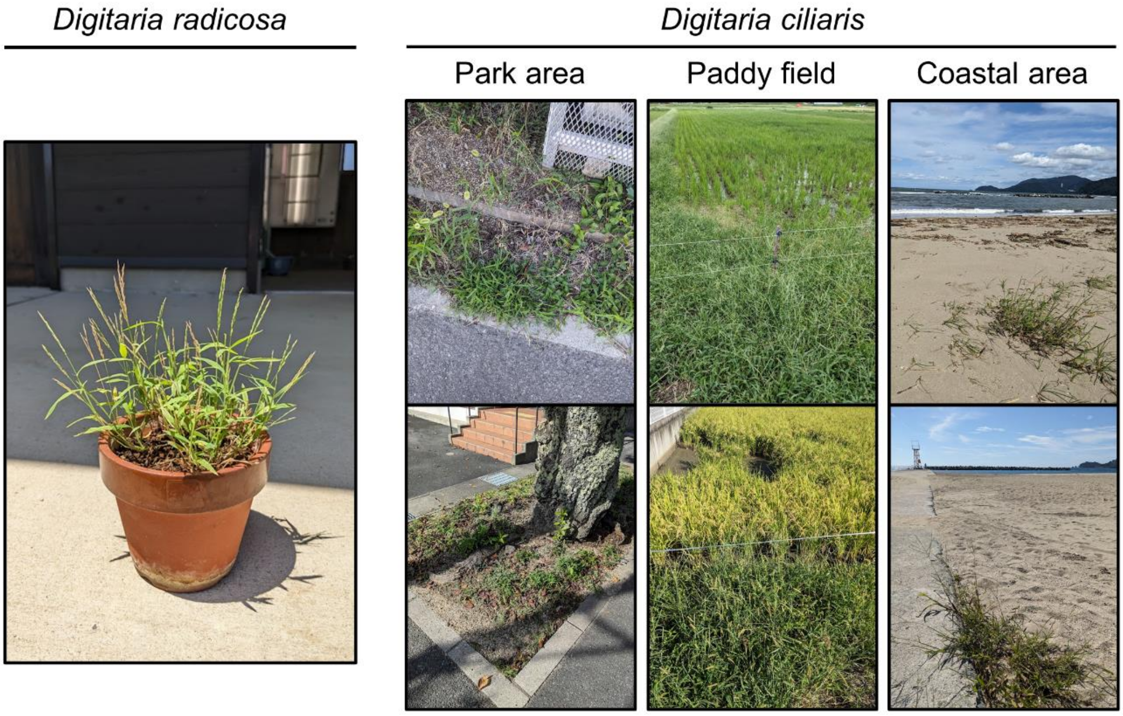


**Figure S1. Example of *Digitaria radicosa* plant and examples of *Digitaria ciliaris* plants growing in three habitats.** Representative example of *D. radicosa* and representative examples of *D. ciliaris* growing in a park area (left), paddy field (middle), and coastal area (right).

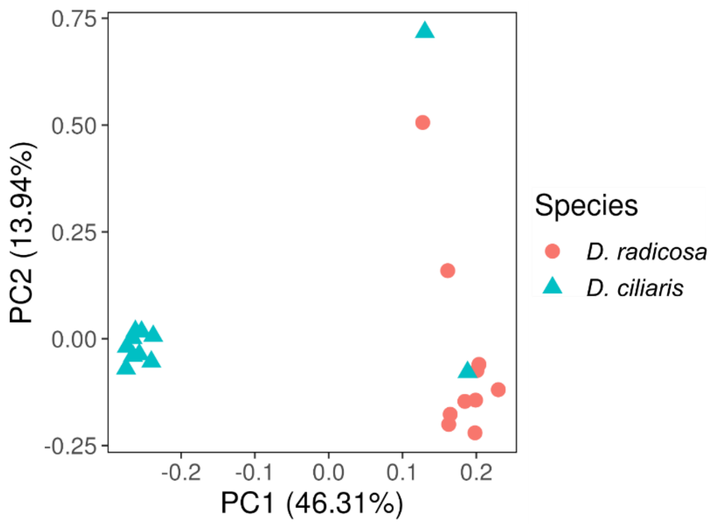


**Figure S2. Principal component analysis of all samples before removing putatively misidentified samples.** Two *D. ciliaris* samples clustered with the *D. radicosa* samples and were suspected to have been misidentified as *D. ciliaris*.

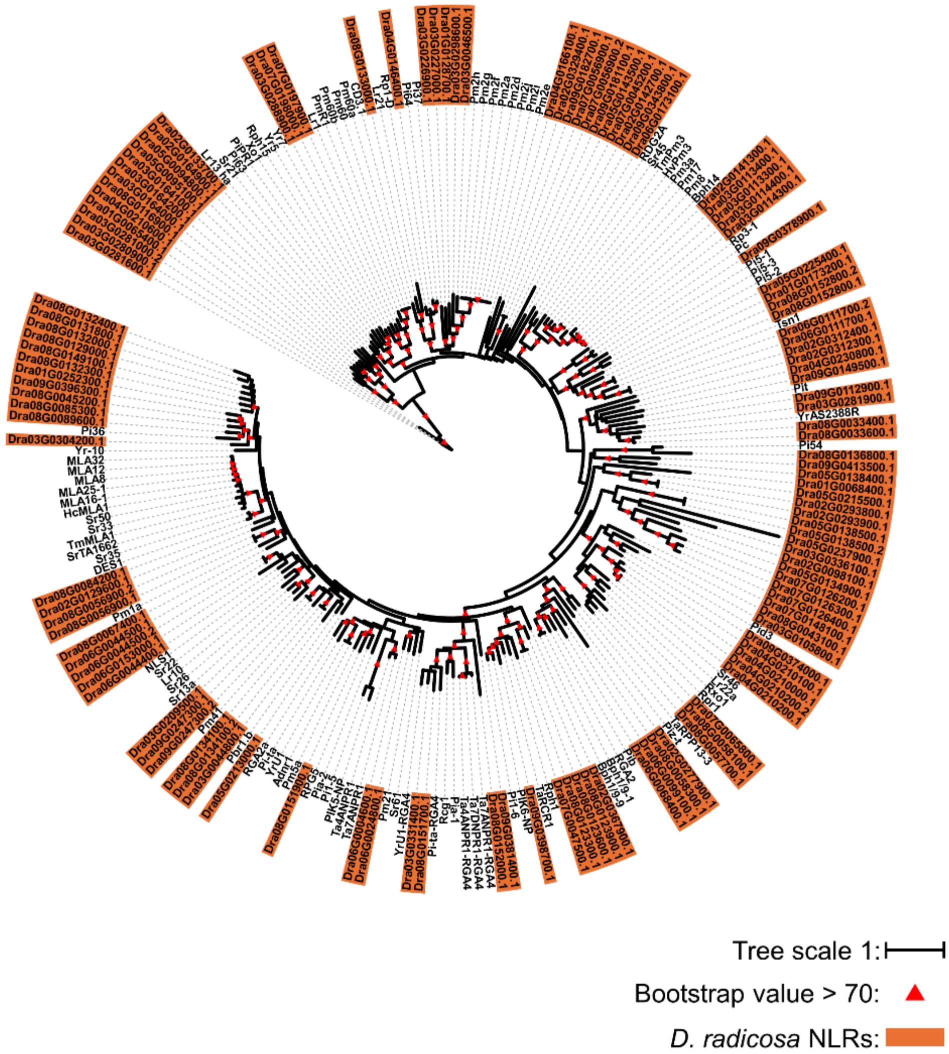


**Figure S3. Phylogenetic tree of identified NLR proteins from *Digitaria radicosa* and functionally validated NLRs from Poaceae species.** Phylogeny of 126 NLR proteins from *D. radicosa* and 100 functionally validated NLR proteins from Poaceae species. The maximum-likelihood phylogenetic tree was generated in RAxML version 8.2.12 with the JTT model using the amino acid sequences of NB-ARC domains. *D. radicosa* NLRs are highlighted in orange. The red arrowheads indicate bootstrap support > 0.7 based on 100 iterations. The scale bar corresponds to the mean number of amino acid substitutions per site on the respective branch.
